# SCARF: Single Cell ATAC-seq and RNA-seq Foundation model

**DOI:** 10.1101/2025.04.07.647689

**Authors:** Guole Liu, Yongbing Zhao, Yingying Zhao, Tianyu Wang, Quanyou Cai, Xiaotao Wang, Ziyi Wen, Lihui Lin, Ge Yang, Jiekai Chen

## Abstract

Recent advances in single-cell multi-omics have provided unprecedented insights into gene regulation by jointly profiling transcriptomic (scRNA-seq) and chromatin accessibility (scATAC-seq) landscapes. However, the inherent heterogeneity and high dimensionality of these multimodal data present significant challenges for effective integration and downstream analysis. Foundation models have demonstrated strong representation learning capabilities for scRNA-seq or scATAC-seq data. So far, however, no model has been specifically developed for the integrative analysis of these two modalities. Here, we introduce SCARF, a <u>s</u>ingle <u>c</u>ell <u>A</u>TAC-seq and <u>R</u>NA-seq <u>f</u>oundation model. SCARF is pre-trained on X-Omics, the largest curated collection of single-cell multi-omics data to date, comprising over 2.7 million cells across multiple tissues and species. The model utilizes a Mamba architecture for efficiently capturing long-context relationships between genes and between accessible regions. Modality-specific and shared features are learned by the model through self-supervised learning and contrastive learning, respectively. SCARF achieves state-of-the-art performance on multiple downstream tasks, including cell representation, cell matching, and cross-omics translation. Furthermore, SCARF enables few-shot cell type annotation, demonstrating strong generalizability across previously unseen datasets. These results highlight the power of foundation models for advancing integrative analysis of single cell multi-omics data, with broad applications in important tasks including cellular characterization, gene or genomic perturbation analysis, and regulation network analysis.

## Introduction

Recent advances in single-cell multi-omics have significantly deepened our understanding of cellular heterogeneity and gene regulation, largely owing to the development of high-resolution transcriptomic and epigenetic profiling techniques^1, 2^. Technologies such as single-cell RNA-seq and ATAC-seq have unveiled the complex interplay between gene expression and chromatin accessibility^3^. However, the inherent heterogeneity and high dimensionality of these datasets pose a major challenge for their effective integration, which is critical to fully utilizing the complementary information they provide.

To overcome these limitations, researchers have increasingly turned to foundation models, deep learning models pre-trained on massive and diverse datasets to extract robust and transferable biological representations. These foundation models then serve as a foundational base for a wide range of downstream tasks^4-7^. In the scRNA-seq field, models such as scBERT^8^ and scGPT^9^ have demonstrated that transformer-based architectures can effectively capture subtle variations in cellular states using tens of millions of single-cell profiles. Likewise, foundation models developed for broader genomic applications, such as DNABERT^10^ for DNA sequence analysis, have showcased the power of pre-training strategies in extracting meaningful biological features.

Current approaches for integrative analysis of single-cell multi-omics data, such as uniPort^11^ and scGLUE^12^, have enabled joint interpretation of scRNA-seq and scATAC-seq datasets. However, these methods face two critical limitations. First, they are predominantly trained and applied on specialized datasets, relying on localized feature harmonization or graph-based alignment techniques. While effective within their original experimental contexts, their generalizability to unseen data remains uncertain due to the lack of cross-dataset biological priors and transferable representations. Second, existing frameworks underutilize the vast repositories of publicly available single-cell data, which now encompass millions of cells across diverse tissues and conditions. By processing each dataset in isolation rather than learning population-scale patterns, current methods are limited in capturing conserved regulatory principles that could strengthen model robustness.

Despite the advances so far, a significant gap remains. Namely, to date, no foundation model has been specifically developed for the integrated analysis of single-cell multi-omics data that simultaneously incorporates both scRNA-seq and scATAC-seq profiles. Although advanced biomedical models like Geneformer^13^, scFoundation^14^ and GeneCompass^15^ have successfully integrated diverse datasets and achieved cross-species analyses, they have not fully addressed the unique challenges posed by single-cell multi-omics data.

Here, we introduce SCARF, a <u>s</u>ingle <u>c</u>ell <u>A</u>TAC-seq and <u>R</u>NA-seq <u>f</u>oundation model. By combining the strengths of both single-cell and general genomic foundation models, SCARF is capable of concurrently capturing transcriptional dynamics and chromatin states. This unified approach not only improves the accuracy of cell type classification and functional annotation but also facilitates the zero-shot matching for unpaired data. In this study, we detail the design and implementation of SCARF, compare its performance with existing state-of-the-art methods in downstream tasks, including cell clustering, cell matching, and cell-type annotation. Our study demonstrates the power of SCARF to advance our understanding of the regulatory mechanisms governing complex cellular processes.

## Results

### SCARF, a <u>s</u>ingle <u>c</u>ell <u>A</u>TAC-seq and <u>R</u>NA-seq <u>f</u>oundation model

We developed SCARF, a foundation model designed to integrate single-cell RNA-seq and ATAC-seq data, addressing the challenges inherent in multi-modal single-cell analysis. We curated X-Omics, a human pre-training corpus comprising over 2.7 million single-cell multiome profiles across diverse tissues, ensuring the model’s capacity for generalization. Leveraging the Mamba architecture^16^, SCARF efficiently learns representations from large-scale, high-dimensional datasets.

In the initial phase of pre-training, SCARF processes RNA and ATAC data separately, converting raw gene expression and chromatin accessibility values into discrete token embeddings. By randomly masking portions of these tokens, the model learns to reconstruct missing signals, capturing both global and context-specific patterns. Subsequently, inspired by the CLIP^17^ (Contrastive Language-Image Pre-training) approach, we introduced a multi-modal fusion layer that aligns the learned RNA and ATAC embeddings into a shared latent space. This integration enhances the model’s ability to capture subtle regulatory signals that might be elusive when analyzing each modality independently (Fig. 1a).

**Figure 1.**
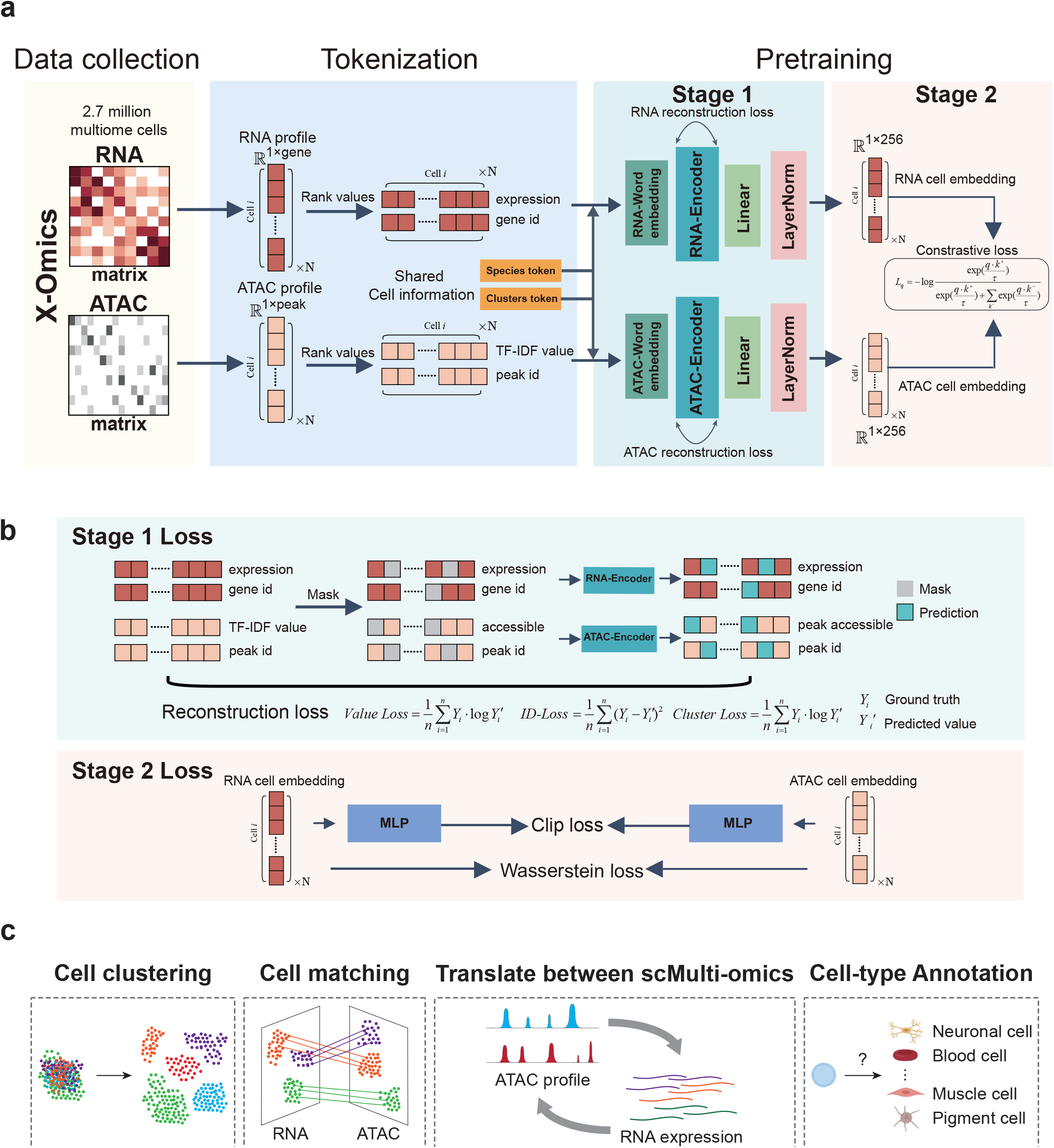
Overview of the SCARF. (a) The workflow of SCARF. Schematic representation of the initial pretraining phase, illustrating how SCARF processes RNA and ATAC data separately to generate discrete token embeddings. The model employs a random masking strategy to learn the reconstruction of missing signals, capturing both global and context-specific patterns. The multi-modal fusion layer aligns the learned RNA and ATAC embeddings into a shared latent space. (b) Illustration of the multi-task loss function used during training. It combines reconstruction objectives, contrastive loss for modality alignment, and clipping-based regularization to reduce outlier impact, ensuring robust learning of single-cell multi-omics representations. (c) Schematic representation of several downstream tasks executed by SCARF, including cell clustering, cell matching, the translation between single-cell multi-omics and cell-type annotation.

During training, we employed a multi-task loss function that combines reconstruction losses to learn modality-specific features, a contrastive loss to align the both modalities, and a clipping-based regularization strategy to mitigate the influence of outliers (Fig. 1b). These components collectively ensure that SCARF learns robust and transferable representations of single-cell multi-omics data. The Mamba architecture’s self-attention and gating mechanisms facilitate the efficient processing of long sequences and sparse signals, maintaining computational feasibility without compromising representational depth.

By integrating scRNA-seq and ATAC-seq data within a unified framework, SCARF enables more precise cell type classification and uncovers novel gene regulatory interactions. This methodology underscores the potential of foundation models to advance the field of single-cell multi-omics, offering deeper insights into complex cellular processes and expediting the discovery of new regulatory mechanisms (Fig. 1c).

### X-Omics, a large-scale and cross-organ pre-training dataset for single-cell Multiome data

To develop SCARF, we established a harmonized multimodal pre-training corpus through systematic integration of single-cell multi-omics datasets capturing paired transcriptomic (RNA-seq) and chromatin accessibility (ATAC-seq) profiles at single-cell resolution from human cells. Our X-Omics resource aggregates 2.7 million high-quality cells derived from 454 publicly available 10x Multiome profiles in the Gene Expression Omnibus (GEO) data repository, spanning 21 tissue types across developmental, homeostatic, and disease contexts (Fig. 2c-d).

**Figure 2.**
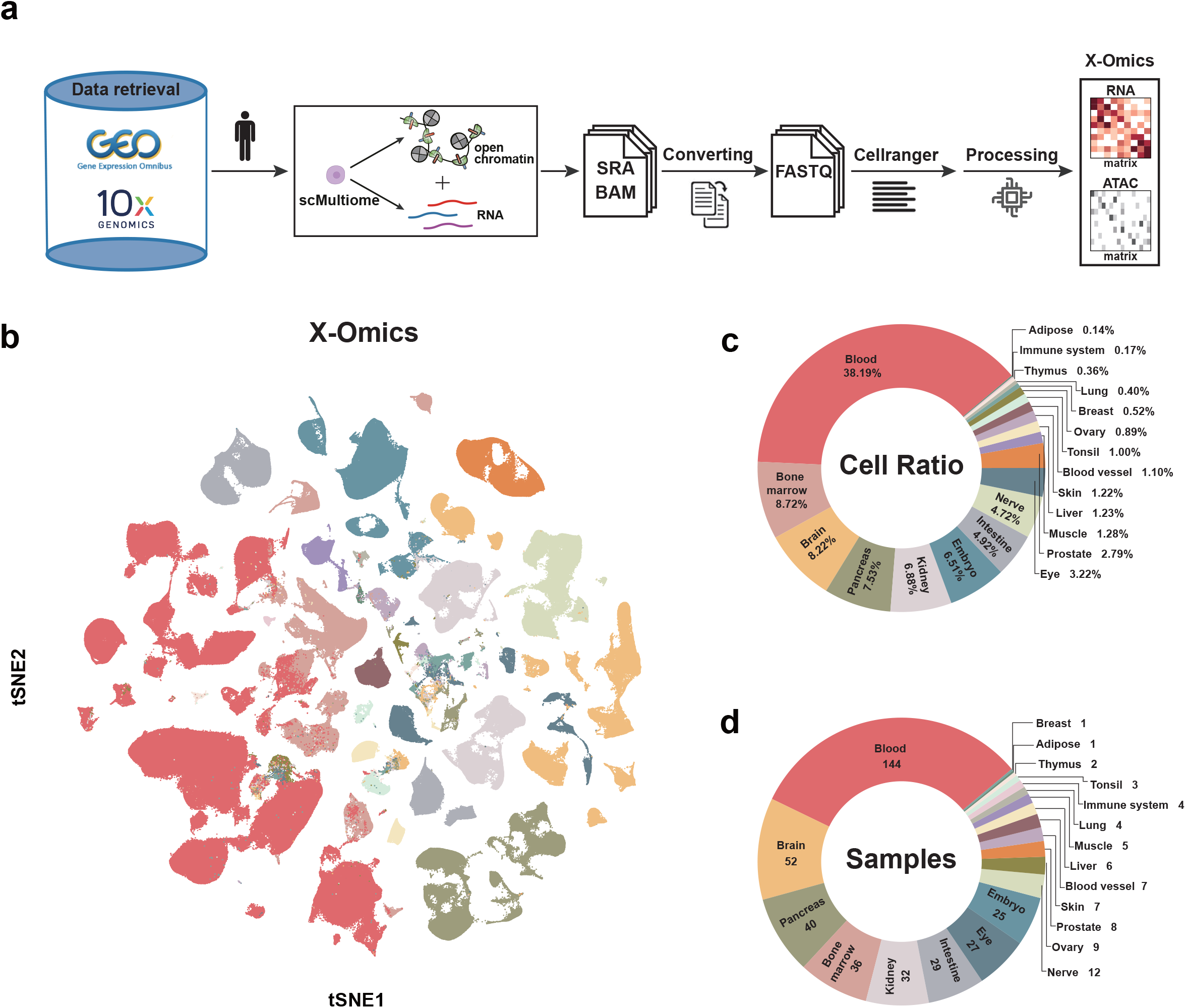
Construction of X-Omics and data overview. (a) Schematic flowchart depicting the methodology used for constructing the X-Omics. (b) t-SNE visualization depicting the distribution of X-Omics cells, classified by tissue types. (c) Proportions of cells corresponding to 21 distinct tissue types. (d) Summary of the number of samples collected for each tissue type.

To ensure consistency across heterogeneous datasets, we implemented a unified processing framework initiating from raw sequencing inputs. Primary SRA files and BAM alignments were systematically reprocessed through standardized workflows involving: (1) FASTQ format conversion, (2) genomic alignment, (3) stringent quality control, (4) data normalization, and (5) generation of cell-by-peak (ATAC) and cell-by-gene (RNA) count matrices (Fig. 2a and Supplementary Fig1, Methods). This comprehensive reprocessing strategy effectively eliminated technical biases inherent in the original 454 datasets, enabling robust integration of epigenetic and transcriptional features.

t-Distributed Stochastic Neighbor Embedding (t-SNE) visualization of X-Omics illustrates the cellular distribution across various tissue types (Fig. 2b). To our knowledge, X-Omics represents the largest curated collection of uniformly processed single-cell Multiome data. This expansive and diverse pre-training resource not only underpins the robust learning of both universal and modality-specific features by SCARF but also provides a critical foundation for downstream applications.

### Zero-shot Cell representation across scRNA-seq and scATAC-seq data

Building on the robust representations learned during pre-training, we next evaluated SCARF’s performance in generating biologically informative cell embeddings from multiome data. We compared SCARF with established methods (scGPT^9^, scFoundation^14^, and scBasset^18^) that rely on a single modality (RNA or ATAC) for representation learning, three multiome datasets obtained from the 10X Genomics website, specifically the hBrain and hPBMC datasets, along with bone marrow mononuclear cells^19^, referred to as the hBMMC dataset.

UMAP plots of cell embeddings derived from RNA, ATAC, and the concatenation of RNA and ATAC embeddings (Fig. 3a) reveal that the concatenated embedding produced by SCARF achieves substantially improved separation of major cell types compared with the single-modality embeddings generated by the alternative models. Quantitative evaluation further supports these observations, with the concatenated RNA□+□ATAC embedding attaining the highest scores across several clustering metrics, including NMI, ARI, AMI, ASW, and homogeneity (Fig. 3b), surpassing the RNA-centric approaches. Overall performance, as summarized by the AvgBio score (Methods) across the three datasets (Fig. 3c), consistently favors the concatenated RNA□+□ATAC embedding. In addition, a heatmap depicting the normalized proportions between clustering categories and true cell types, along with recall scores (Fig. 3d), confirms that SCARF’s integrated approach yields the strongest concordance with ground truth cell type annotations.

**Figure 3.**
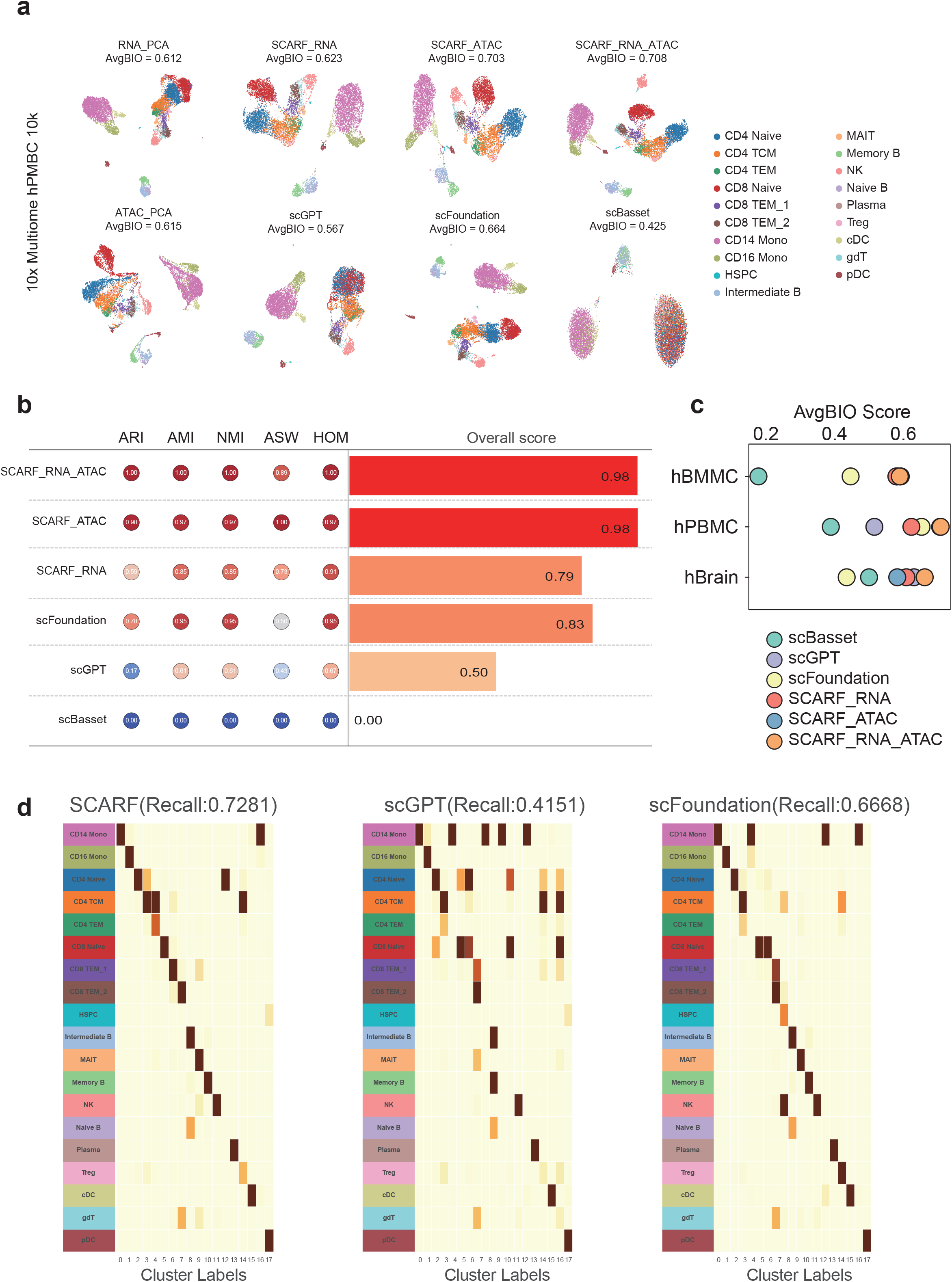
Benchmarks on zero-shot cell clustering performance. (a) UMAP visualizations of 10X Multiome PBMCs data are shown, based on embeddings from RNA PCA, ATAC PCA, scGPT, scFoundation, scBasset, and SCARF using RNA, ATAC, and concatenated RNA + ATAC, colored by cell type. (b) A comparison of clustering performance metrics (ARI, AMI, NMI, ASW, and HOM) for scGPT, scFoundation, scBasset, and SCARF using RNA, ATAC, and concatenated RNA + ATAC embeddings, along with an overall score for each method. (c) AvgBIO Scores for scGPT, scFoundation, scBasset, and SCARF using RNA, ATAC, and concatenated RNA + ATAC embeddings across hBMMC, hPBMC, and hBrain datasets. (d) A heatmap illustrating the normalized proportions between clustering categories and true cell types, along with recall scores for SCARF, scGPT, and scFoundation methods.

These results indicate that concatenating RNA-seq and ATAC-seq embeddings via SCARF enhances cell type discrimination and clustering accuracy, underscoring the promise of foundation models in advancing single-cell multiome analyses.

### Zero-shot cell matching across scRNA-seq and scATAC-seq data

To assess the performance of modality matching between RNA and ATAC, we conducted UMAP visualization of the latent embeddings from SCARF and several competing methods (scGLUE^12^, Seurat^20^, Harmony^21^, LIGER^22^, uniPort^11^, and scVI^23^). The UMAP results demonstrated that cells from different modalities clustered closely in the shared latent space, highlighting the effective matching and integration achieved by SCARF (Fig. 4a). To quantitatively benchmark modality matching, we used the matching probability score, which measures the accuracy of cell pairings (Fig. 4b). Our method outperformed the others, achieving the highest score on the 10X Multiome data of human peripheral blood mononuclear cells (PBMCs). Additionally, we employed the Fraction of Samples Closer Than the True Match (FOSCTTM) score to evaluate soft matching (Fig. 4c). SCARF yielded the lowest score among all methods, with lower values indicating better matching, thus demonstrating superior modality alignment.

**Figure 4.**
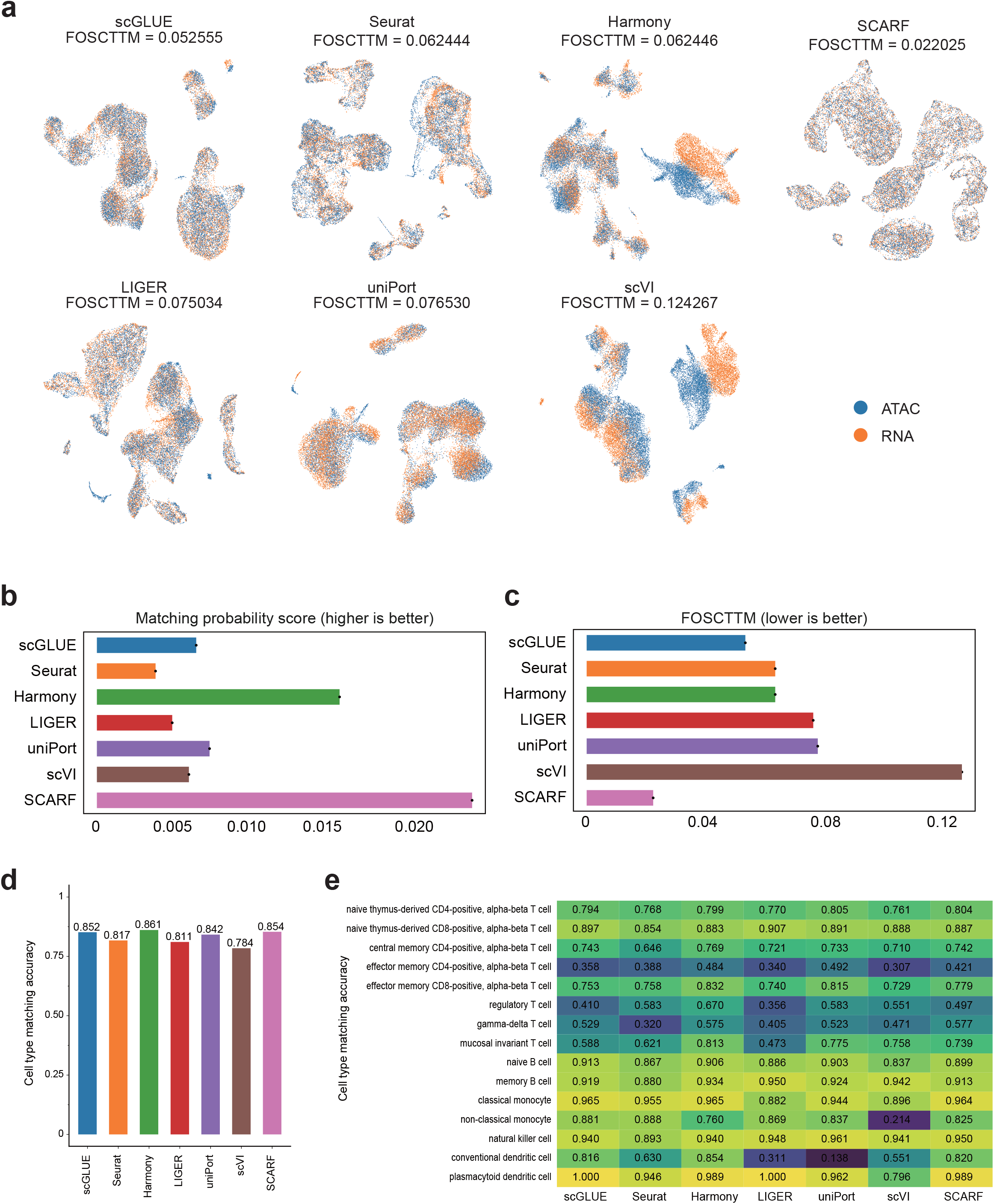
Benchmarks on zero-shot cell matching performance. (a) UMAP visualizations of integrated cell embeddings for 10X Multiome PBMCs data, colored by modalities. (b-c) Benchmarking cell matching using the matching probability score (b) and FOSCTTM score (c). Higher scores indicate better performance for the matching probability score, while lower scores are preferred for the FOSCTTM score. (d-e) Benchmarking the accuracy of matching cells to the correct cell type, shown overall in (d) and for each individual cell type in €.

To further analyze whether the mismatched cells were actually matched to cells of identical cell types, we calculated cell type matching accuracy, where a match is defined as pairing cells of the same type (Fig 4d, e). We found that SCARF achieved a high cell type matching score of 0.854, ranking second only to Harmony. Upon examining the scores for each cell type, we observed that our method achieved scores above 0.8 for 9 out of 15 cell types, with the remaining lower scores corresponding to related cell types.

### Zero-shot cross-omics translation between scRNA-seq and scATAC-seq data

To further assess the capacity of SCARF capacity to learn modality-invariant representations, we evaluated its ability to predict RNA-based profiles from ATAC-seq inputs by mapping query data to reference embedding (Fig. 5a, Methods). Specifically, we provided the model with chromatin accessibility data and examined its predictions of gene expression. SCARF-predicted RNA embeddings closely resemble the actual RNA embeddings, displaying comparable clustering of major immune cell subtypes(Fig. 5b). Quantitative metrics including ARI, NMI, and HOM demonstrate that the predicted RNA embeddings preserve critical cell type distinctions at levels approaching those observed with real RNA data.

**Figure 5.**
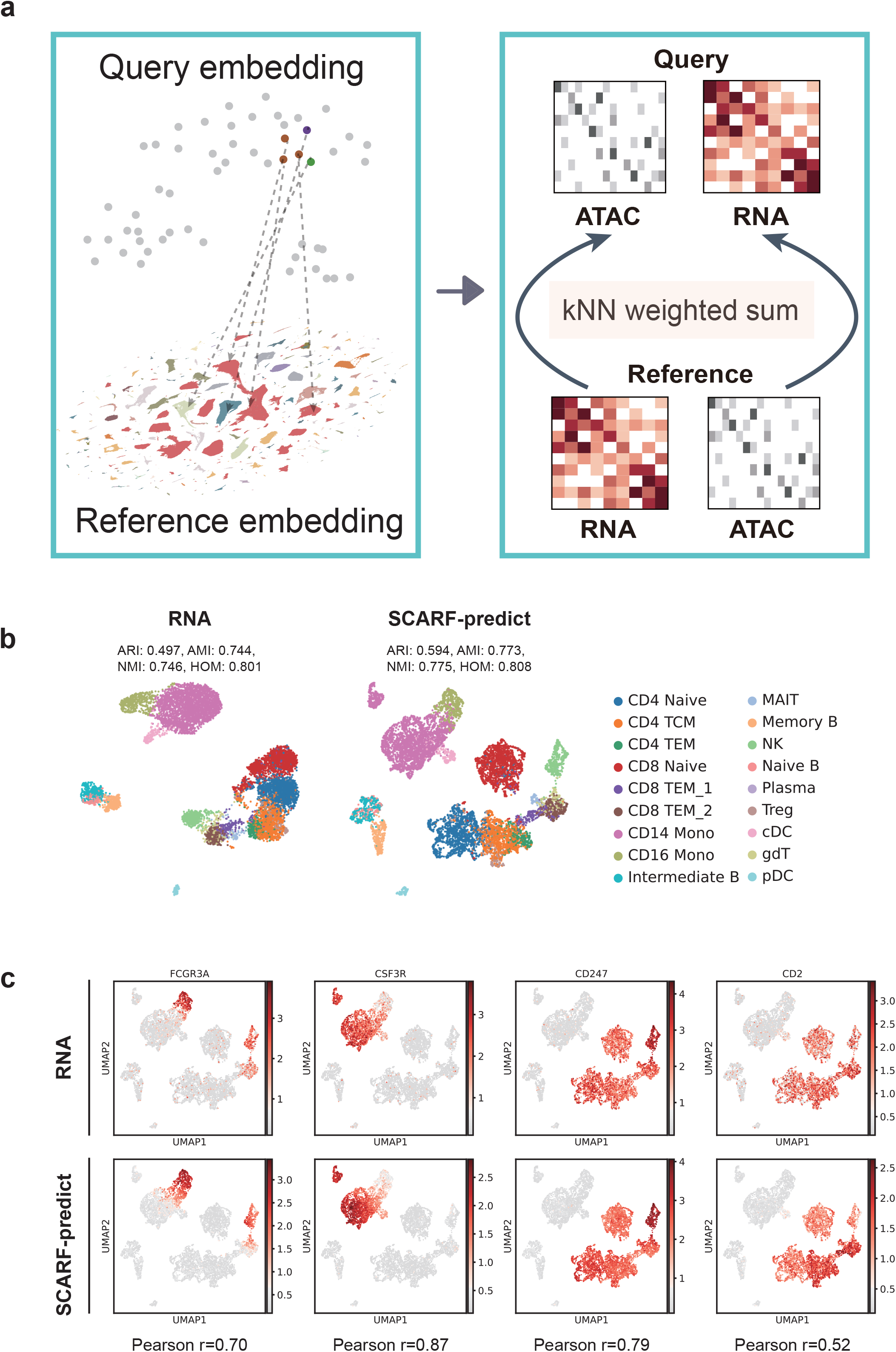
Zero-shot translation between single-cell multi-omics. (a) An illustration of cross-omics prediction is provided. (b) UMAP visualizations show the actual RNA data and the predicted RNA data from SCARF, along with clustering performance metrics (ARI, AMI, NMI, ASW, and HOM), colored by cell type. (c) UMAP projections of the actual and predicted values of cell type marker genes, such as FCGR3A, CSP3R, CD247, and CD2, along with Pearson correlation coefficients for several genes between the true and predicted values.

Moreover, we investigated the correspondence between predicted and actual gene expression values (Fig. 5c). UMAP projections of representative genes, such as FCGR3A, CSP3R, and CD247, revealed a high degree of correlation (Pearson r = 0.70-0.87) between the SCARF-predicted and ground truth measured expression patterns. Notably, the spatial distributions of these predicted gene expressions were closely aligned with the underlying cell clusters, indicating that SCARF effectively integrates chromatin accessibility information to infer transcriptional states.

Together, these findings suggest that SCARF learns robust cross-modality representations, enabling accurate reconstruction of RNA-seq profiles from ATAC-seq inputs. This capability highlights the potential utility of foundation models in single-cell multiome analyses, particularly in settings where one modality is missing or less accessible.

### Few-shot cell type annotation

To fine-tune the pre-trained SCARF for cell type annotation, we employed a logistic regression classifier, using the cell embeddings output from SCARF as input and predicting cell type classifications. The model was trained on a reference dataset with expert annotations (10% of the cells) and then used to predict the cell types in the held-out query dataset (90% of the cells). We conducted extensive experiments across multiple datasets to evaluate the performance of SCARF in cell type annotation (Fig. 6a).

**Figure 6.**
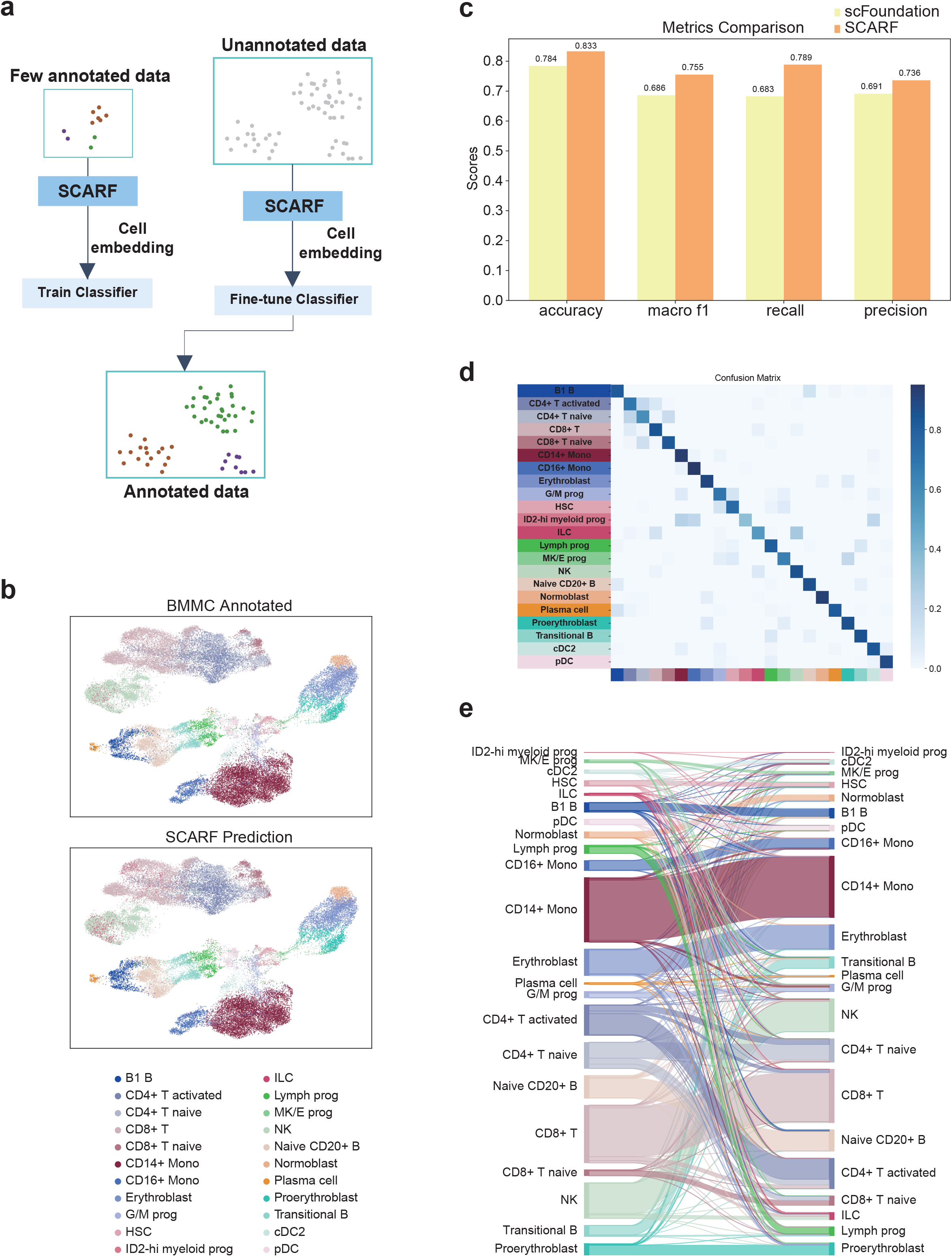
Benchmarks on cell-type annotation performance. (a) An illustration of fine-tuning the SCARF model using a small number of annotated cells to achieve cell-type annotation tasks. (b) UMAP visualizations of true labels and predicted labels for BMMC data, colored by cell type. (c) Bar charts comparing cell annotation evaluation metrics (accuracy, macro F1, recall, and precision) for scFoundation and SCARF. (d) A heatmap displaying the confusion matrix information for predicted results across different cell types. (e) A Sankey plot showing the relationship between true labels and predicted labels for cell types.

The predicted cell clusters exhibited a strong correspondence with annotated cell types, with distinct immune populations remaining well separated (Fig. 6b). Evaluation of annotation performance (Fig. 6c) revealed accuracy, macro F1, recall, and precision scores above 0.75, better than single modality foundation model scFoundation^14^. This alignment is further supported by the confusion matrix, which shows high agreement between the model’s predictions and ground truth labels, with minimal misclassification across cell types (Fig. 6d). Sankey diagram (Fig. 6e) visualizes the correspondence between predicted labels and reference annotations, illustrating how SCARF effectively preserves the underlying cellular hierarchy across various lineages, including lymphoid and myeloid compartments.

These findings indicate that the learned representations of SCARF enable accurate few-shot cell type annotation, even in the absence of large scaled data fine-tuning. This capability holds significant promise for accelerating the characterization of complex single-cell datasets, particularly in resource-limited or exploratory settings.

## Discussion

Our study introduces SCARF, a foundation model that integrates scRNA-seq and ATAC-seq data through the efficient Mamba architecture, providing a robust framework for multi-modal single cell analysis. Our findings indicate that SCARF not only captures high-quality transferable representations from a large and diverse pre-training corpus but also performs exceptionally well in downstream applications, including cell representation, modality alignment, cross-omics translation, and cell type annotation.

The improved cell clustering achieved through concatenated RNA and ATAC embeddings illustrates the benefits of a multi-modal-integrated approach over single-modality methods. Moreover, the excellent performance in aligning multi-modal signals and correcting batch effects confirms that SCARF effectively harmonizes heterogeneous datasets while preserving essential biological information. The model’s capacity to accurately infer gene expression profiles from chromatin accessibility data further reinforces its utility, particularly in settings where single data modality is limited or unavailable.

The results in cell type annotation demonstrate the broad applicability of SCARF, suggesting that the representations learned during pretraining are highly generalizable across diverse biological contexts. These results open new avenues for future work, including the incorporation of additional omics layers, such as proteomics or spatial transcriptomics, and the potential use of the model in clinical settings to support rapid diagnostic and prognostic evaluations. We plan to refine SCARF by investigating more advanced fusion strategies and expanding its ability to process even larger, more complex datasets. Moreover, adapting the model for real-time analysis in high-throughput settings could further accelerate translational research and enhance our understanding of cellular heterogeneity and gene regulatory networks.

## Methods

### Collection of the X-Omics

We obtained publicly available 10x Genomics single-cell multiome (ATAC + Gene Expression) datasets from the Gene Expression Omnibus (GEO) repository. Sample annotations were manually curated based on GEO metadata entries and corroborated with information from associated publications.

Raw sequencing files were acquired through multiple repositories including the European Bioinformatics Institute (EBI), Amazon Web Services (AWS), and China National Center for Bioinformation (CNCB). For samples with available sequence read archives (SRA), we performed format conversion using fasterq-dump (v2.10.9) with default parameters to generate paired-end FASTQ files. When both BAM and SRA files were available, we prioritized the use of BAM files and converted them to FASTQ format using Cell Ranger ARC’s bamtofastq utility (v2.0.2), following the recommended protocols by 10X Genomics. File provenance was verified through read length analysis during format conversion.

All sequencing data were aligned to reference genomes (GRCh38) using Cell Ranger ARC (v2.0.2) with default mapping parameters. The pipeline simultaneously processed chromatin accessibility and gene expression data while performing cell calling, peak calling, and generating feature-barcode matrices through integrated analysis of both modalities.

### Data quality control

Following initial processing with Cell Ranger ARC (v2.0.2), we implemented a two-tiered quality control workflow. First, sample-level filtering was performed to exclude low-quality datasets using thresholds of median UMI counts > 80 for RNA and median high-quality fragments > 250 for ATAC.

At the cellular resolution, we applied modality-specific quality filters:

- For RNA modality:

1. Doublets identified via Scrublet^24^ (v0.2.3) with default parameters
2. Minimum 200 detected genes (nFeature_RNA)
3. Minimum 1,000 UMIs (nCount_RNA)
4. Maximum mitochondrial content ratio of 0.2 (calculated from MT-gene subset)

- For ATAC modality:

1. Doublets identified via ArchR^25^ (v1.0.2) with default parameters
2. Minimum 1,000 high-quality fragments (nFragments)
3. Minimum Transcriptional Start Site (TSS) enrichment score of 4

### Single-cell RNA-seq processing

We selected protein-coding genes as features and constructed a cell-by-gene UMI count matrix.

### Single-cell ATAC-seq processing

To ensure consistency of features across samples, we created a reference peak set consisting of 1,743,872 peaks, which were aggregated from 254 ATAC-seq datasets. A two-step noise reduction protocol was implemented:

1. Sample-specific denoising: Peaks were independently called for each sample using ArchR^25^ (v1.0.2), with removal of fragments outside these regions.
2. Reference-based quantification: Filtered fragments were quantified against the reference peaks to generate a unified peak-by-cell matrix.

The matrix was binarized and transformed using TF-IDF, with the inverse document frequency (IDF) calculated across all aggregated single-cell Multiome ATAC datasets to capture the overall feature prevalence.

### SCARF tokenization, architecture, and pre-training

#### SCARF tokenization

The SCARF is pre-trained with a total of 2.7 million single-cell multiome profiles. The scRNA-seq corpus contains human cells with a total gene count of 19,365 protein-coding genes. For scRNA-seq data, we use a ranked tokenization strategy similar to GeneCompass^15^. Firstly, the count sum for non-zero genes of each cell are normalized to 10,000 and log1p transformation is used for gene counts. We then randomly sample 100,000 cells and calculate the median of each gene using the non-zero expression genes. Then we normalized the expression of each gene using its corresponding median to obtain the relative expression level of each gene in the whole scRNA-seq corpus. Besides the gene id and gene expression, we add contextual tokens <CLS> to the input data for each cell. The <CLS> token encodes the semantic features that can represent the whole single cell and can be directly used for cell classification tasks. Finally, each cell can be denoted as following:

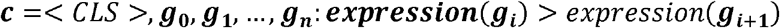

where *expression*(*g*_i_) is a function that returns the expression of gene *i*.

#### SCARF architecture

SCARF is designed as a unified framework for integrating single-cell RNA-seq and ATAC-seq data. The model leverages a modified mamba architecture and a series of specialized modules to learn robust representations from high-dimensional, multi-modal single-cell data. Its design comprises several components: input representation, positional encoding, modality-specific encoding, fusion and modality generation modules, and projection layers for downstream tasks.

#### Input Representation and Positional Encoding

Each cell is represented by two sets of raw data: gene expression values (scRNA-seq) and chromatin accessibility signals (ATAC-seq). These raw measurements are preprocessed and discretized into tokens(as described in the Tokenization section).

Each token is then mapped into a high-dimensional continuous space using embedding layers. Positional encodings are added to the token embeddings to capture sequential or spatial context. Formally, if *E*(*x*) denotes the embedding of a token *x* and *P*(*i*) is the positional encoding at positio*n i*, the final input representation is given by:

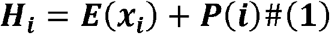

#### Modality-Specific Encoding

Separate encoder modules are employed for the two modalities. For RNA data, the RNA encoder applies self-attention to capture intra-modality dependencies, while ATAC data are processed via the ATAC encoder. During pre-training, a fraction of the tokens is masked, and the model learns to reconstruct the missing values. The reconstruction loss for a given modality is computed as:

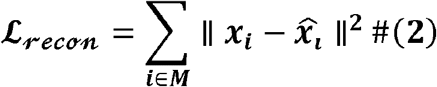

where *M* denotes the set of masked positions, *x*_*i*_ is the original token value, and 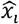 is the model’s prediction.

#### Fusion and Cross-Modality Integration

After modality-specific processing, cell-level representations ([CLS] token) are extracted and projected into a common latent space via dedicated fully connected layers (denoted as *f*_*RNA*_ and *f*_*ATAC*_). These projected embeddings are then normalized:

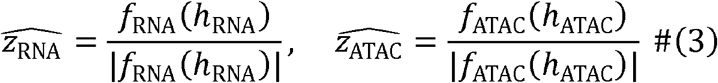

Inspired by the CLIP framework, a contrastive loss is applied to align the RNA and ATAC embeddings. The inter-modality contrastive loss is defined as:

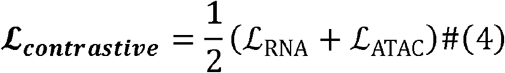

where the loss for the RNA modality is computed using cross-entropy:

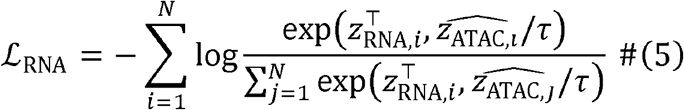

The overall training loss is a weighted sum of several components:

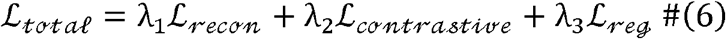

where λ_*i*_ are scalar weights balancing the contribution of each term, and *L*_*reg*_ includes regularization terms (such as clipping-based penalties) to stabilize training.

In summary, the SCARF architecture integrates a modified mamba backbone with modality-specific encoders, a fusion module inspired by CLIP. By combining these components with a multi-task loss function and a dynamic queue for contrastive learning, SCARF effectively captures both shared and modality-specific features from single-cell multiome data.

#### SCARF training

The model was trained using the AdamW optimizer with a weight decay of 0.001. A cosine learning rate schedule was applied, with the learning rate warming up for 1,000 steps before decaying to a minimum value. The maximum learning rate was set to 5×10−^4^. The batch size was 32, with gradient accumulation set to 1 step to accommodate memory constraints. Gradient clipping was applied to prevent exploding gradients. Training was conducted for four epochs using mixed-precision (BF16/FP16) computation on 8 NVIDIA GPUs in a distributed setting with torch.distributed.launch. Checkpoints were saved at regular intervals, and model selection was based on validation loss. To improve contrastive learning stability, a queue mechanism was implemented to maintain a dynamic set of RNA and ATAC embeddings, updating at each training step to enhance negative sampling efficiency.

### Datasets, evaluation and analysis for the cell matching downstream task

We used 10x demo multiome data of human peripheral blood mononuclear cells (PBMCs), which consists of 12,073 cells across 15 distinct cell types, to perform modality matching. The embeddings obtained from SCARF, which are derived from cells of different modalities, were optimized for better modality alignment using the optimal transport Sinkhorn algorithm. To quantitatively evaluate modality matching, we employed two metrics: the matching probability score and the Fraction of Samples Closer Than the True Match (FOSCTTM) score. The matching probability score measures the accuracy of correctly paired cells, where higher scores indicate better matching between modalities. On the other hand, the FOSCTTM score evaluates soft matching by calculating the proportion of samples that are closer to the true match compared to random pairings, with lower FOSCTTM scores suggesting better alignment of modalities. We employed the cell type matching accuracy to assess the correctness of cell type matching across different modalities. A cell pair is deemed correctly matched if both cells belong to the same cell type in two modalities. Higher accuracy scores reflect better alignment of cells from different modalities, indicating superior performance in correctly identifying matched cell types.

The matching probability score is computed as:

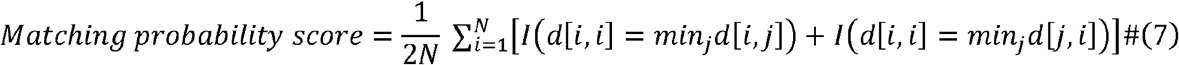

The FOSCTTM score is computed as:

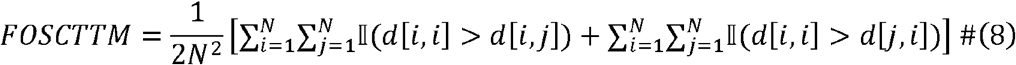

### Datasets, evaluation and analysis for the cross-omics translation downstream task

The cross-omics prediction method is zero-shot, leveraging precomputed reference RNA and ATAC embeddings along with query data embeddings to achieve efficient conversion from RNA to ATAC. Initially, the RNA and ATAC embedding matrices and cell names are loaded from the pre-generated reference dataset. Using the RNA embedding from the query data, a fast KNN search (with k=10) is performed on the reference RNA embeddings via the fastKnn function (which employs cosine distance as the similarity metric and adjusts neighbor and index construction parameters accordingly) to identify the most similar reference cell for each query cell. Subsequently, the distances for each query cell’s neighbors are transformed into weights using a softmax function, assigning greater influence to the more similar neighbors. Based on these weights, the method reconstructs the query cell’s ATAC expression profile by iterating over the corresponding neighbor indices, retrieving the ATAC expression data from the reference dataset, and computing a weighted average of the neighbors’ ATAC expressions, thereby generating a new ATAC expression vector for each query cell.

For the quantitative evaluation of the translated profiles in terms of preserving cell heterogeneity, we employed four metrics to assess the clustering results: adjusted Rand index (ARI), adjusted mutual information (AMI), normalized mutual information (NMI), and homogeneity (HOM). In our analysis, we considered a set of translated profiles comprising *N* cells categorized into cell types *T* = {*T*1,… …,*Tn*} and cluster labels *P* = {*P*1,… …, *Pm*}, which were derived using the Leiden algorithm. Here, *a*_*i*_ and *b*_*j*_ denote the counts of cells corresponding to labels *T*_*i*_ and *P*_*j*_, respectively, while *n*_*ij*_ represents the number of cells that fall into the overlap between *T*_*i*_ and *P*_*j*_.

The ARI metric accounts for chance agreement and is computed as an adjustment of the Rand index (RI) through the following formula:

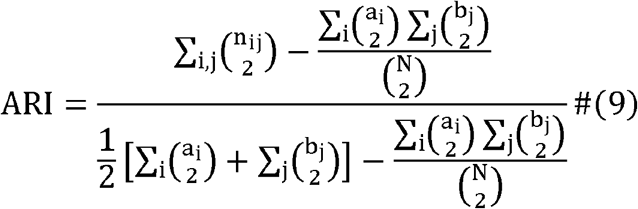

NMI is a normalized variant of mutual information (MI), which could be represented as:

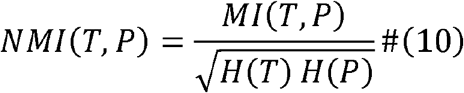

where *MI*(·) is the mutual information:

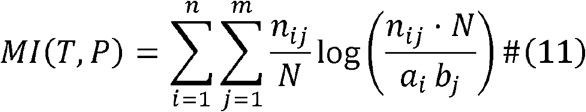

and *H*(·) represents the entropy.:

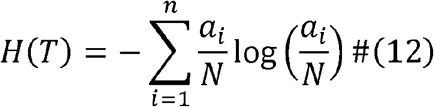

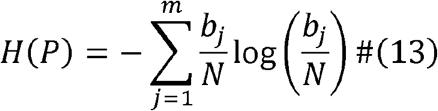

AMI further considers the chance agreement and is written as:

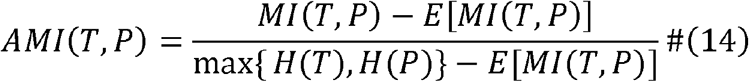

HOM provides a reference for the purity of cell types within each cluster, calculated as follows:

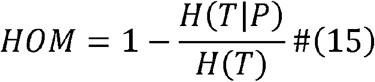

where *H*(*X*|*Y*) means the conditioned entropy of *X* under the condition that *Y* is given.

In our analysis of ATAC chromatin accessibility, we employ the area under the receiver operating characteristic curve (AUROC) as a metric to assess the quality of our predictions. For RNA expression, which we treat as a continuous variable, we utilize Pearson’s correlation coefficient to evaluate the accuracy of our RNA expression predictions and gene activity scores. Both scatterplots and density scatterplots are annotated with the corresponding Pearson’s correlation values. Given that single-cell RNA sequencing (scRNA-seq) analyses, such as clustering and dimensionality reduction, are primarily conducted on log-transformed counts, all correlation computations and visual representations are performed in log space. In our approach, we consider each gene or peak in each individual cell as a distinct observation or prediction for the purposes of both correlation analysis and AUROC evaluation.

### Datasets, evaluation and analysis for the cell representation downstream task

For this evaluation, we use two scRNA-seq datasets: the hPBMC dataset and the hBMMC dataset. Compared to single-modality models such as scGPT^9^ and scBassat^26^.

We evaluate the representation capability of SCARF through clustering tasks, focusing on how well the learned representations group similar cells. Clustering performance is quantified using metrics such as Adjusted Rand Index (ARI), Average Silhouette Width (ASW), Normalized Mutual Information (NMI), and the comprehensive metric Avgbio, which integrates biological relevance across these measures.

Avgbio integrates multiple clustering metrics to evaluate the biological relevance of the clustering results.

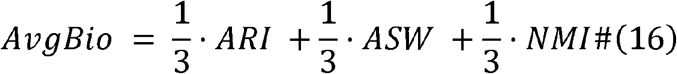

### Datasets, evaluation and analysis for the cell type annotation downstream task

For the cell type annotation task, we utilized the same dataset as in the previous tasks, ensuring consistency in evaluation across different methods. The datasets include multi-modal single-cell profiles, such as RNA-seq and ATAC-seq, derived from public repositories 10X website. We compared the performance of foundation models, such as scFoundation. Foundation models leverage large-scale pre-training to generalize across tasks and datasets.

For evaluating the performance of cell type annotation models, we used the following classification metrics. Their formulas are provided below: Accuracy measures the proportion of correctly classified instances among all instances.

Precision measures the proportion of correctly predicted instances for a specific class out of all instances predicted as that class. It is computed for each cell type and averaged (micro, macro, or weighted).

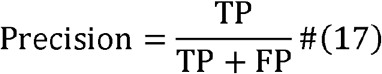

Recall evaluates the proportion of correctly predicted instances for a specific class out of all actual instances of that class.

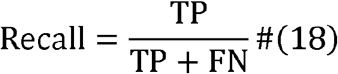

The F1-score is the harmonic mean of precision and recall, providing a single metric to balance the trade-off between them.

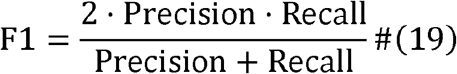

## Data availability

The X-Omics data were sourced from the GEO database, with a cutoff date of February 23, 2024. The hPBMC and hBrain datasets can be accessed through the 10X Genomics website. Additionally, the hBMMC dataset was collected from the NCBI Gene Expression Omnibus (GEO) under accession number GSE194122, which can be found at the following link: [https://www.ncbi.nlm.nih.gov/geo/query/acc.cgi?acc=GSE194122].

## Code availability

SCARF is implemented in Python based on the PyTorch framework. The source code for reproduction is available at https://github.com/JiekaiLab/scarf and https://github.com/cbmi-group/scarf.

## Acknowledgements

This work was supported by the National Natural Science Foundation of China (grant no. 32225012 to JK.C., grant no. 92354307 to G.Y., grant no. 32200662 to LH.L., grant no. 32401262 to GL.L.), National Key Research and Development Program of China (grant no. 2023YFF1204701 to JK.C., grant no. 2024YFF0729202 to G.Y., grant no. 2024YFA1802300 to YB.Z.), the Strategic Priority Research Program of the Chinese Academy of Sciences (grant no. XDA0460305 to G.Y.), the Fundamental Research Funds for the Central Universities (grant no. E3E45201×2 to G.Y.), Science and Technology Planning Project of Guangdong Province (grant no. 2023QN10×323, 2023ZT10Y154, 2023B1212060050, 2023B1212120009 to YB.Z.) and Health@InnoHK Program launched by Innovation Technology Commission of the Hong Kong SAR, China.

## Author Contributions

JK.C., G.Y., YB.Z. and LH.L conceived the study and supervised the project. GL.L designed SCARF frameworks and contributed to code. TY.W. and YY.Z curated and processed data in X-Omics. TY.W. performed the evaluation of cell matching. QY.C. performed the evaluation of cell clustering. QY.C. and ZY.W performed the evaluation of cell-type annotation. QY.C., YY.Z and XT.W performed the evaluation of translate between scMulti-omics. QY.C. and YY.Z wrote the manuscript.

## Supplementary tables and figures

**Supplementary Figure 1.**
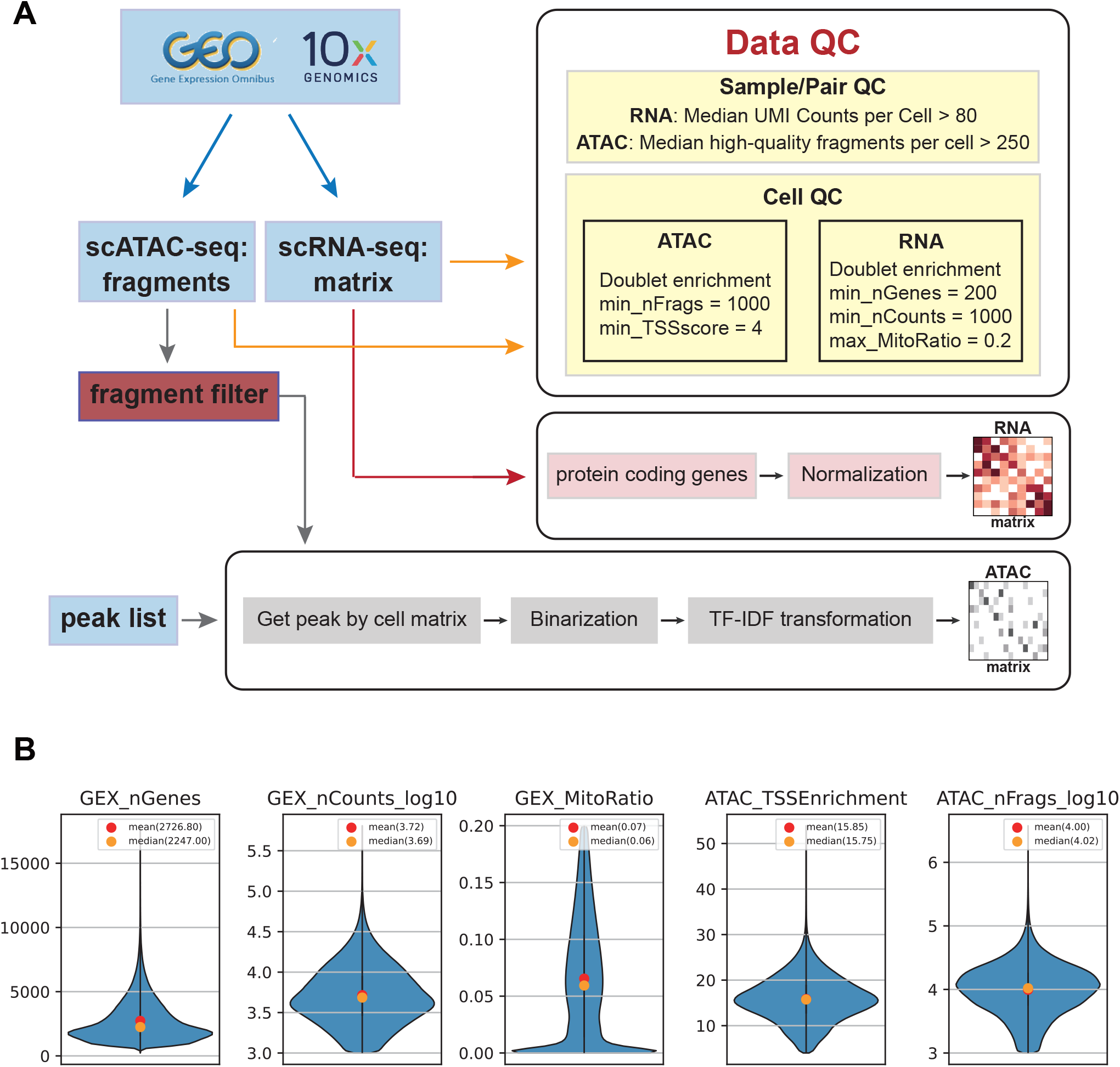
Overview of data processing workflow. (a) Workflow illustrating the steps of quality control and the construction of RNA and ATAC matrices. (b) Quality control metrics for cells, including number of genes, UMI counts, mitochondrial ratio, TSS enrichment score, and number of fragments.

**Supplementary Figure 2.**
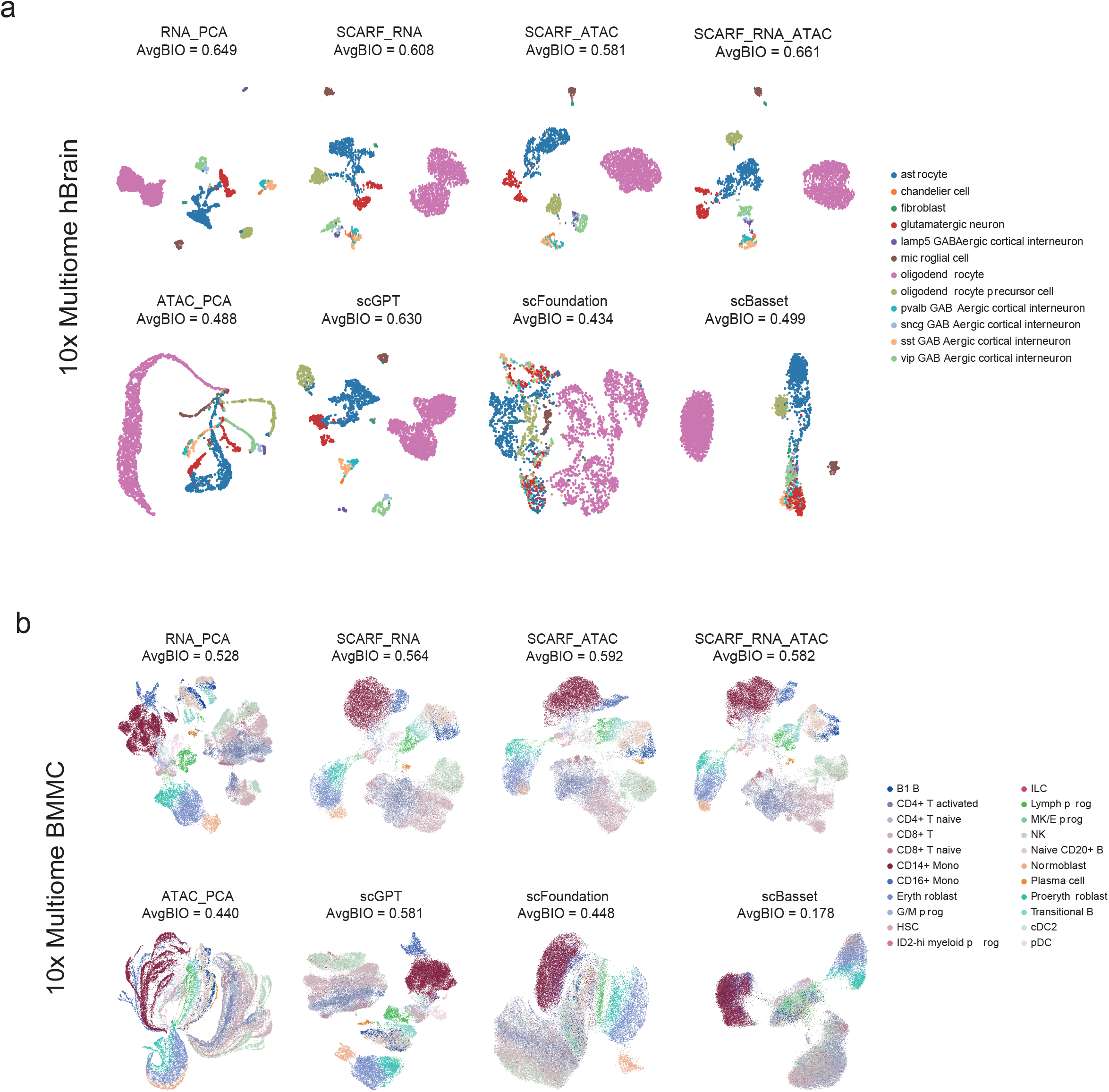
Performance Evaluation of Cell Clustering Methods. (a-b) UMAP visualizations of 10X Multiome data for hBrain (a) and BMMC (b), colored by cell type, with AvgBIO scores annotated on the plots for different methods.

